# An RNAi screen in *C. elegans* for genes that play a role in secretion and cleavage of VAPB MSP domain

**DOI:** 10.1101/2021.01.02.425092

**Authors:** Hala Zein-Sabatto, Jim Collawn, Chenbei Chang, Michael A. Miller

## Abstract

VAPB (VPR-1 in *C. elegans*) is a type-II ER transmembrane protein whose N-terminal Major Sperm Protein domain (MSPd) is cleaved and secreted. Mutations in the MSPd of human VAPB cause impaired secretion and are associated with Amyotrophic Lateral Sclerosis (ALS). In *C. elegans*, the secreted MSPd signals non-cell-autonomously to regulate striated muscle mitochondrial morphology and gonad development. As VAPB/VPR-1 does not have a signal peptide and its MSPd extends into the cytosol, it is unclear how the protein is proteolytically cleaved and secreted. To identify genes that are involved in VPR-1 cleavage and unconventional secretion, we performed an RNA interference (RNAi) screen in *C. elegans*. Worms null for *vpr-1* are sterile and have striated muscle mitochondrial abnormalities. These defects can be rescued by *vpr-1* expression in the neurons, germline, or intestine, implying that these three tissues share a common machinery to cleave and secrete the MSPd. Examination of shared gene expression in these tissues revealed a list of 422 genes, which we targeted with RNAi. *vpr-1* null worms expressing *vpr-1* from intestine were used in the screen, and the brood size of these worms after RNAi knockdown was scored. Disruption of factors involved in VPR-1 MSPd processing and/or secretion should revert fertility phenotypes in these worms. We identified many genes that induce compromised fertility when knocked down in these but not wild type worms, including a V-SNARE, several proteasome components, stress response molecules, and mitochondrial genes. Our screen thus identified many potential players involved in MSPd processing and/or secretion.

**Summary:** The MSP domain (MSPd) of a type-II ER transmembrane protein called VAPB is cleaved and secreted to function as a non-cell-autonomous signal. The topology of VAPB positions MSPd in the cytosol. It is thus unclear how MSPd is cleaved from VAPB and released extracellularly. Using *C. elegans*, we screened 422 genes by RNAi to identify potential candidates regulating MSPd cleaving and secretion. We identified the Golgi v-SNARE YKT-6 and several components of the 20S and 19S proteasome that may mediate MSPd trafficking and cleaving, respectively. These results have promising implications in advancing our understanding of MSPd signaling.

## Introduction

Human VAPB is a member of the VAP (<u>V</u>AMP/synaptobrevin <u>a</u>ssociated <u>p</u>roteins) family of proteins. VAPs are endoplasmic reticulum (ER) membrane proteins with roles in membrane trafficking, ER unfolded protein response, and lipid transport at intracellular membrane contact sites (1, 2). Three domains make up the VAPB protein: a cytosolic N-terminal MSP domain (MSPd), a central coiled-coil domain, and a C-terminal ER-anchoring transmembrane region (1, 3–5). The MSPd with an immunoglobin-like β-sheet structure is named for its homology with a well-characterized nematode protein known as the Major Sperm Protein, or MSP (1, 3, 6). Nematode MSP is an abundant sperm cytoskeletal protein required for sperm motility (7, 8). In *C. elegans*, it is secreted from sperm in an unknown fashion to regulate oocyte maturation (9–11). The MSPd from VAPB is cleaved and can also be released to the extracellular space via an unconventional pathway in a cell type-specific fashion (12–15). However, the mechanism by which the MSPd is proteolytically processed and unconventionally secreted is not yet understood.

In humans, the MSPd has been detected in blood and cerebral spinal fluid (CSF) and implicated in fatal neurodegenerative diseases, such as Amyotrophic Lateral Sclerosis (ALS) and spinal muscular atrophy (SMA) (12, 16). An amino acid substitution of proline 56 with serine (P56S) in the MSPd of VAPB prevents MSPd secretion and precisely segregates with familial cases of ALS (12, 17–20). In *C. elegans* and *Drosophila*, the MSPd secreted from neurons binds to receptors on striated muscles and in gonadal cells to modulate mitochondria positioning to the myofilaments and gonad development, respectively (12–15, 21). *C. elegans* null for *vpr-1*, the VAPB homolog, are sterile and have body wall muscle mitochondrial abnormalities (15, 21). These phenotypes can be rescued by expression of *vpr-1* solely in the germline, or intestine, or the nervous system (15). These data thus support a model of an endocrine, non-cell-autonomous role for the MSPd in signaling.

To identify genes essential for VAPB/VPR-1 MSPd processing and unconventional secretion, we conducted an RNAi screen in *C. elegans*. Based on the rationale that the cleavage and the unconventional secretion machinery is shared among the three tissues whose expression of *vpr-1* can rescue the null phenotypes, we assembled a list of 422 genes which are expressed commonly in the germline (22), intestine (23), and nervous system (24). RNAi was performed to knockdown these genes in a rescue line expressing *vpr-1* driven by an intestinal promoter, *ges-1p* (*ges-1p::vpr-1*). Brood size was scored to identify genes that decreased the fertility of the rescue line but did not affect the fertility of wild type worms. Many genes without any previously reported function in regulating protein processing or secretion have been uncovered in our study.

## Methods and Materials

### *C. elegans* Strains

*C. elegans* strains were propagated at 20°C and fed with NA22 *E.coli* (25), unless otherwise indicated. N2 Bristol was the wild type strain used in the RNAi screen. The transgenic line used in this screen *(ges-1p::vpr-1)* is VC1478 *vpr-1(tm1411)/hT2 [bli-4(e937) let-?(q782) qIs48] (I; II)* transgenically expressing extrachromosomal arrays of *ges-1p::vpr-1* (originally made in Cottee et al., 2017).

### RNAi Screen

RNAi knock down of genes was performed by the feeding method (26). HT115(DE3) bacterial feeding strains were obtained from the Ahringer library (27). PCR and sequencing (UAB Heflin Center for Genomic Sciences) were used to confirm that bacterial strains contained the correct clones. Each RNAi bacterial feeding strain was grown to feed parallel plates of N2 and the transgenic *ges-1p::vpr-1* worms (Figure 1). Five L4 staged worms of each genotype were maintained on respective RNAi plates at 25°C. Progeny was counted after 7 days on RNAi plates at 25°C. RNAi knockdown was replicated three times for *ykt-6, rpn-1, rpn-10*, and*pbs-2*. RNAi constructs for *pas-4* and empty L4440 backbone were used as positive and negative controls, respectively. Student’s t-test statistical analysis was conducted using Prism software. Pie chart was made with Microsoft Excel.

**Figure 1:**
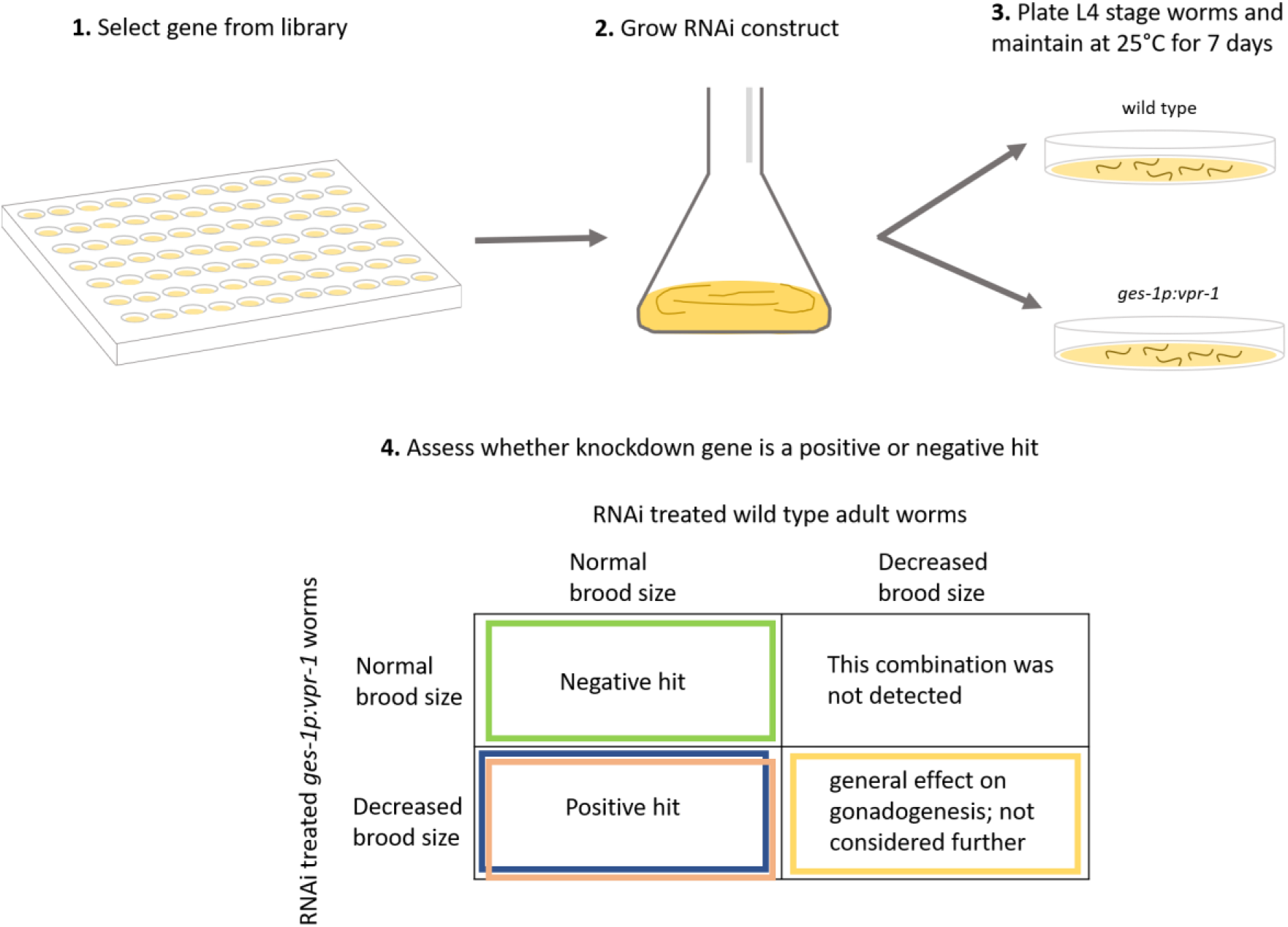
RNAi Screen Paradigm to identify potential effectors of MSPd cleaving and secretion. Colors highlighting hits coordinate with list of genes in Table S1.

### Data Availability

Any worm strain or reagent used in this screen is available upon written request to the corresponding author. All data sets and results from this screen are included in this report.

## Results and Discussion

*C. elegans* muscle mitochondria and gonadogenesis are regulated non-cell-autonomously by the cleavage and secretion of MSPd from VAPB/VPR-1 (12–15, 21). However, genes that play a role in the cleaving or secretion of the MSPd have yet to be identified. This screen aimed to take an unbiased approach to identify potential players in these processes. Identifying such players can have implications for neurodegenerative diseases like ALS that has been linked to the lack of MSPd circulation in human blood and cerebral spinal fluid (12, 16).

The design of this screen is based on previous work from our lab. We showed *vpr-1* expression in neuronal, intestinal, or germline cells is sufficient to rescue the *vpr-1* null phenotypes of sterility and muscle mitochondrial abnormalities in *C. elegans* (15, 21). This result suggested that these three cell types may share a common mechanism that detaches the MSPd from VPR-1 and releases the MSPd extracellularly into the pseudocoelom to be accessed by the body wall muscles and somatic gonad. Therefore, gene expression data sets of *C. elegans* nervous system (24), intestine (23), and germline (22) were compared. These data sets revealed 422 overlapping genes that became the targets of this screen (Table S1).

*C. elegans* null for *vpr-1* are maternal effect sterile (15). Of the three strongest rescue lines, intestinal transgenic expression of *vpr-1* in a *vpr-1* null background *(ges-1p::vpr-1)* was chosen to be used in this screen. A strength of using the *ges-1p::vpr-1* line is that it provides a non-redundant system of VAPB/VPR-1 MSPd cleaving and secretion that is RNAi-sensitive. Transgenic expression exclusively in the germline would require integration into the genome to avoid silencing and neurons are resistant to the effect of RNAi (15, 28). *vpr-1* null worms are maternal effect sterile, but the majority of *ges-1p::vpr-1* worms are fertile (15). Taken together, this screen was designed to knock down a gene that plays a role in VAPB/VPR-1 MSPd cleaving or secretion in order to decrease non-cell-autonomous MSPd signaling such that the otherwise fertile *ges-1p::vpr-1* line reverts back to sterility.

Of the 422 selected genes, 71 candidates were not found in the Ahringer RNAi library or did not grow in culture (Table S1). RNAi constructs of the remaining genes were fed in parallel to wild type N2 Bristol and transgenic *ges-1::vpr-1* worms (Figure 1). Any gene whose RNAi knockdown resulted in a decrease of the N2 brood size was not considered a hit because of its direct implication in fertility or gonadogenesis in a wild type background (Figure 1; Table S1). First-tier hits were arbitrarily categorized as RNAi knockdown genes that did not affect the brood size of the N2 worms while the *ges-1p::vpr-1* plates had a count of 20 or less live progeny (Figure 1; Table S1). The number of live *ges-1p::vpr-1* progeny between 20-50 worms after RNAi knockdown constituted the second tier hits. The screen was not designed to identify any gene, such as a negative regulator, that when knocked down enhanced the fertility of the rescued *ges-1p::vpr-1* line.

First and second tier hits were categorized based on their known functions (Figure 2). A portion of these hits (18%) could not be classified or remain uncharacterized. One-third of the hits are involved in RNA/DNA binding. These genes alongside the 4% comprising nucleoporin genes may be upstream players that regulate the transcription, mRNA processing, and mRNA transport of proteins responsible for VAPB/VPR-1 MSPd cleaving or secretion. 13% are involved in cell division and could not be explained with regards to VAPB/VPR-1 MSPd cleaving and secretion. 11% are mitochondrial proteins of which three genes encode mitochondrial ribosomal proteins. These proteins may regulate expression of mitochondria outer membrane proteins that can interact with VAPB/VPR-1 and influence the conformation or accessibility of VAPB/VPR-1 for efficient cleavage or secretion.

**Figure 2:**
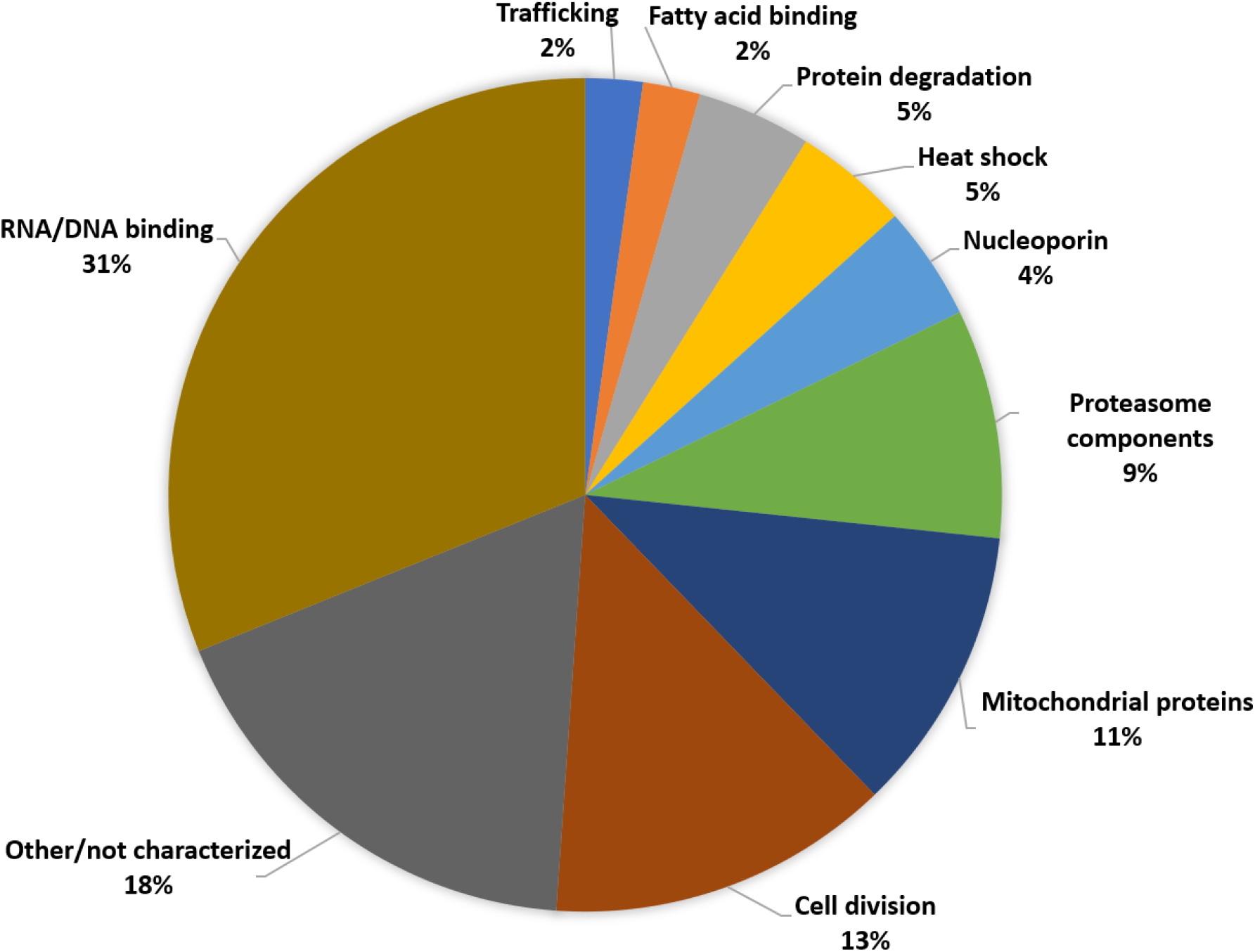
Pie chart depicting the various gene categories identified in this RNAi screen. Genes from the first and second tier of hits were categorized based of off their description on Wormbase.

In the protein degradation category, *ubxn-6* is a member of the ubiquitin regulatory X (UBX) domain proteins and is the *C. elegans* homolog for UBXD-1 in humans (29). UBXD-1 functions as one of several adaptor proteins to an AAA (<u>A</u>TPase <u>a</u>ssociated with various <u>a</u>ctivities) ATPase known as p97/VCP, or CDC-48 (30). One of two CDC-48 isoforms in *C. elegans* was among the 422 genes screened (C06A1.1) but it did not rank in the top two tiers (Table S1). p97/VCP/CDC-48 has different functions based on the binding of various adaptor proteins (31). 97/VCP/CDC-48 binds UBXD1 to mediate ubiquitinated protein trafficking into endolysosomes as well as the formation of ERGIC-53 positive vesicles between the ER-Golgi and potentially the plasma membrane (32, 33). Interestingly, mutations in p97/VCP/CDC-48 that negatively affect UBXD1 binding have been found to segregate with cases of familial ALS, as is the case with the P56S mutation in VAPB that prevents MSPd cleavage and secretion (12, 17, 32, 34, 35). Whether VAPB/VPR-1 MSPd is being trafficked or processed by p97/UBXD1 needs to be further investigated.

Of the 18 first-tier hits, five genes have been previously reported to play a role in protein trafficking or processing (Table 1). Knock down of the v-SNARE *ykt-6* or proteasome components *rpn-1, rpn-10*, and *pbs-2* significantly decreased the brood size of *ges-1p::vpr-1* rescue line, but did not affect N2 fertility (Figure 3). How these candidate genes may be playing a role in VABP/VPR-1 MSPd cleaving or secretion is further discussed below:

**Table 1:**
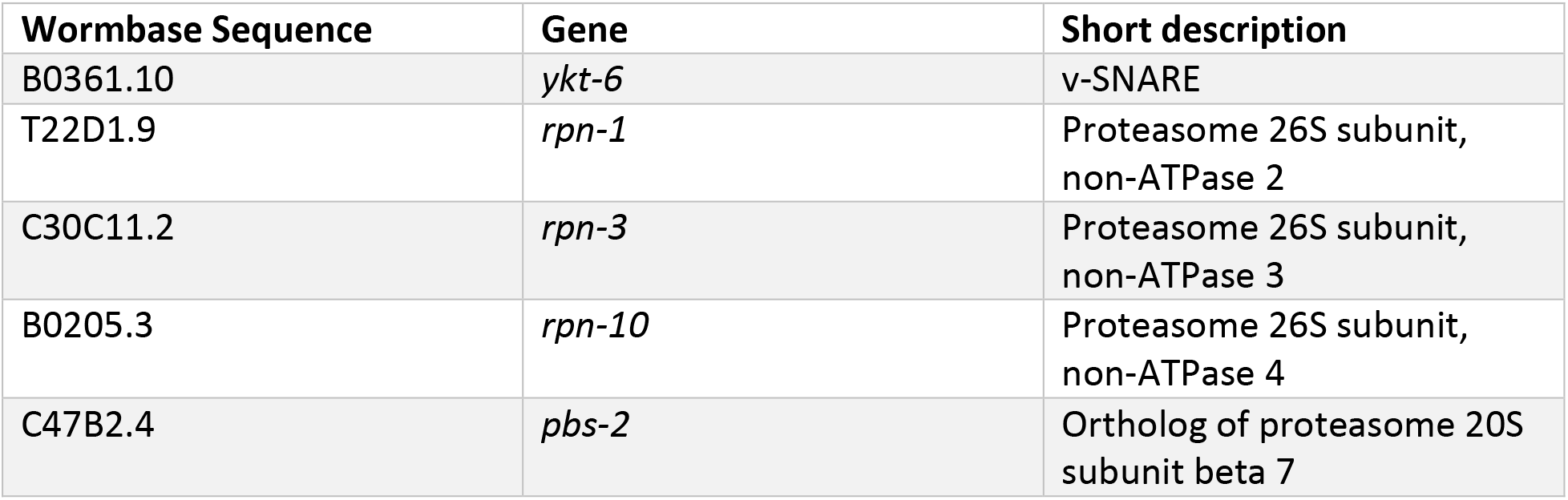
Five selected candidate genes from the top-tier list.

**Figure 3:**
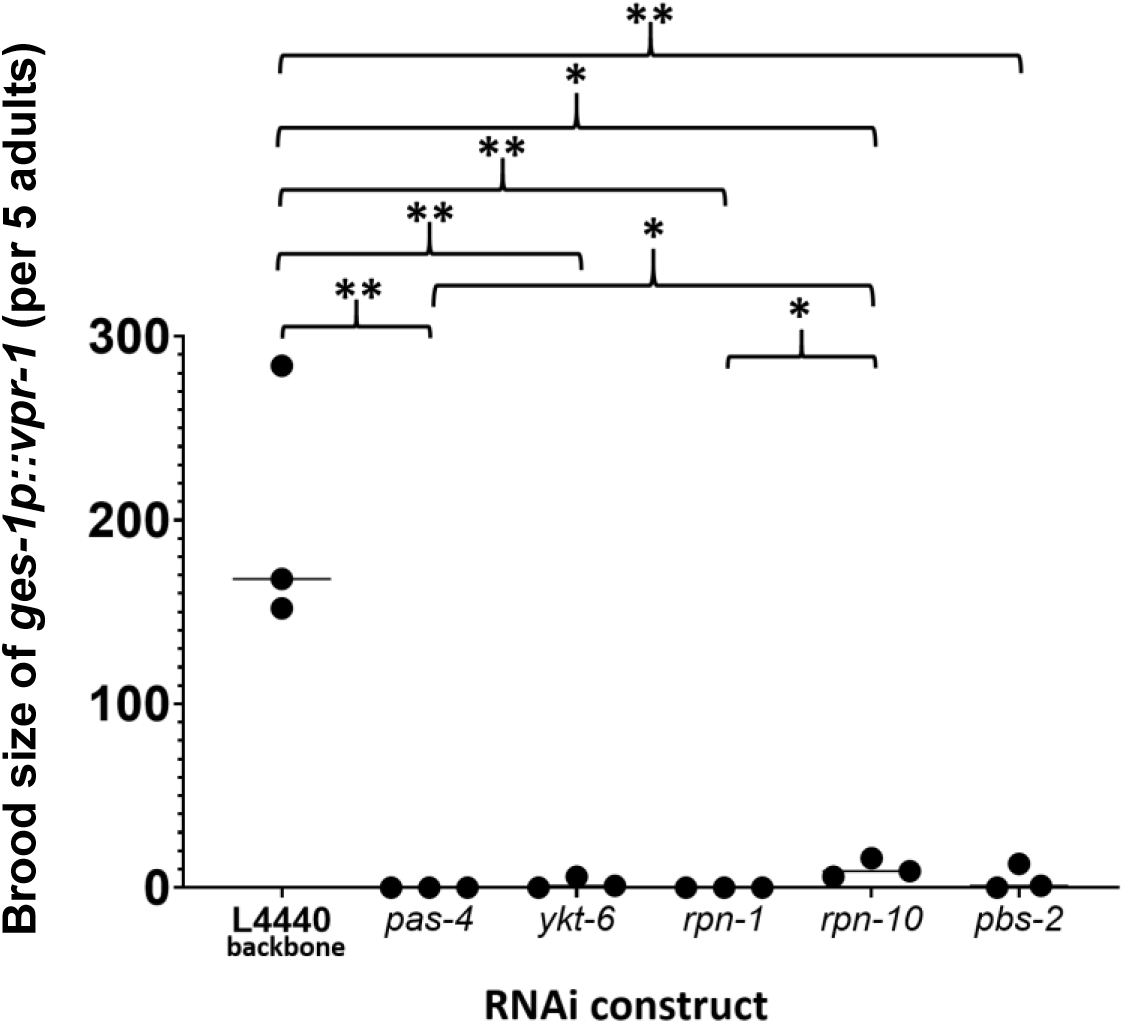
Brood sizes of *ges-1p::vpr-1* decreased after RNAi knockdown of *ykt-6, rpn-1, rpn-10*, and *pbs-2*. Five L4 *ges-1p::vpr-1 C. elegans* were fed RNAi constructs of select gene for 7 days at 25°C. *ges-1p::vpr-1 C. elegans* grown on empty RNAi backbone L4440 produced an average of 200 progeny. The brood size decreased when *ykt-6, rpn-1, rpn-10*, or *pbs-2* were knocked down in *ges-1p::vpr-1 C. elegans*. Knock down of these four genes did not affect N2 brood sizes (not shown). Knock down of *pas-4* was used as an RNAi control since it caused sterility and lethality in both N2 and *ges-1p::vpr-1 C. elegans*. (*p>0.01 and **p>0.001, Student’s t-test)

### A SNARE gene: ykt-6

SNAREs, or soluble N-ethylmaleimide-sensitive factor (NSF) attachment protein receptors, play a crucial role in mediating membrane fusion between vesicles and cellular compartments of the secretory pathway. YKT-6 is a vesicle SNARE, or R/v-SNARE protein, that is further classified as a longin, or long vesicle-associated membrane protein (VAMP) (36). This classification is attributed to its N-terminal profilin-like domain, known as a longin domain (LD), that helps regulate YKT-6 binding (36). Unlike most SNAREs, YKT-6 lacks a transmembrane domain (37, 38). Instead, YKT-6 is anchored into the membrane of various organelles by lipid modification. For example, to anchor into the Golgi membrane, a lipid moiety is attached to the C-terminal CAAX motif by an irreversible post-translational modification, farnesylation (37–39), as well as a reversible post-translation modification, palmitoylation (39, 40). Farnesylation itself is insufficient for YKT-6 to insert into the membrane, as most YKT-6 is present as a depalmitoylated, farnesylated, cytosolic protein (37–39, 41–43). Cytosolic YKT-6 assumes a closed conformation in which the N-terminal LD folds back and binds the SNARE motif to tuck away and shield the hydrophobic farnesyl anchor (44, 45). This conformation maintains YKT-6 in a stable, soluble state and serves to autoinhibit the SNARE motif from binding to and negatively affecting membrane bound SNAREs from forming complexes between vesicles and target membranes (36, 39, 41, 44). This accessible pool of cytosolic YKT-6 may be the reason that YKT-6 is a multifaceted SNARE that functions in various intracellular trafficking routes and is found bound to the Golgi, endosomes, exosomes, lysosomes-autophagosomes, and vacuole membranes in yeast (37, 38, 41–43, 46–52).

Due to the promiscuous nature of YKT-6, it is difficult to explain how YKT-6 might be involved in the cleavage or secretion of VAPB/VPR-1 MSPd. One possibility is that the MSPd is not cleaved from VAPB/VPR-1 until after it reaches the plasma membrane via YKT-6 positive vesicles. While VAPB/VPR-1 does not contain a signal peptide found in conventionally secreted proteins, it does localize to the ER membrane facing the cytosol (4). We suggest that this localization may allow VAPB/VPR-1 to be passively or actively incorporated into COPII vesicles headed for the Golgi, the main site of active bound YKT-6. Transport between the ER and Golgi may be influenced by *ubxn*-6, as discussed above. Once reaching the Golgi, YKT-6 may facilitate further sorting and trafficking of VAPB/VPR-1 through distinct secretory pathways. RNAi knockdown of Ykt-6 in *Drosophila* and human cells revealed that Ykt-6 plays a role specifically in the secretion of Wnt proteins through exosomes (53). VAPB/VPR-1 may be trafficked through a similar exosomal route in a YKT-6-dependent manner. The depletion of YKT-6 in combination with Syb/VAMP3 also resulted in an accumulation of post-Golgi vesicles, indicating that YKT-6 plays an additional role in plasma membrane fusion (50). In neurons, one of the cell types known to secrete the VAPB/VPR-1 MSPd, YKT-6 was reported to localize to unknown membranous compartments, and not the Golgi, in a punctate fashion(15, 41). Therefore, VAPB/VPR-1 may be trafficked by YKT-6 positive vesicles, whether exosomes or otherwise, to the plasma membrane to have the MSPd cleaved before extracellular release. This hypothesis is consistent with our observation of full length VAPB/VPR-1 protein localization at the basolateral membrane of *C. elegans* intestinal cells (54). Further studies to assess a direct relationship between VAPB/VPR-1 and YKT-6 positive vesicles are needed to test this model.

### Proteasome components: pbs-2, rpn-1, rpn-3, and rpn-10

Proteasomes are diverse and have been implicated in the degradation of misfolded or excess protein in order to maintain cellular homeostasis (55). The 20S proteasome has a barrel-like structure with an internal proteolytic core that can function independently or in complex with a regulatory cap referred to as the 19S proteasome to make up the 26S proteasome (56, 57).

The 19S proteasome binds, deubiquitinates, and mediates the translocation of ubiquitinated proteins through the 20S proteasome (58). Two top tier hits from this screen, *rpn-1* and *rpn-3*, encode for two non-ATPase subunits of the 19S proteasome base and lid, respectively (56, 58). Another positive hit, *rpn-10*, encodes for the subunit that links 19S proteasome lid and base, contains two polyubiquitin binding motifs, and functions as one of several ubiquitin receptors (58, 59). These ubiquitin receptors are non-redundant and recognize specific substrates. For example, deleting RPN-10 specifically results in a feminization phenotype in *C. elegans* hermaphrodites due to the accumulation of TRA-2 proteins (60). When knocking down other components of the 19S base from the list of 422 genes (*rpt-1, rpt-3*, and *rpt-5*), the N2 brood size was also affected indicating that these genes may play a broader role in *C. elegans* development. Knocking down other subunits of the 19S lid, such as *rpn-5* and *rpn-12*, had no effect on N2 brood size, but did not decrease the brood size in *ges-1p::vpr-1* as drastically as when knocking down *rpn-1*, *rpn-3*, or *rpn-10*.

Proteolysis, or the breakdown of peptide bonds, occurs in the 20S proteasome. The 20S proteasome is comprised of a stack of two outer rings of α-type subunits and two inner rings of β-type subunits (56). The two inner β-rings encompass the chymotrypsin-like, trypsin-like, and caspase-like proteolytic sites (56). PBS-2 in *C. elegans* is the *β2* subunit, one of three β-subunits that make up these proteolytic sites in the 20S proteasome (56). Knock down of *pbs-2* with RNAi had no effect on wild type *C. elegans* brood size, but rendered the *ges-1::vpr-1* worms sterile (Figure 3). The other two proteolytic subunits, *β1* and *β5*, were not among the 422 genes screened here. Other β-type subunits,*pbs-3,pbs-4,pbs-6*, and *pbs-7*, and α-type subunit *pas-6* were included in the list of 422 candidates, but knockdown of these genes, unlike *pbs-2*, decreased the wild type brood size (Table S1). This indicates that PBS-2, unlike the other 20S proteasome subunits, may play a specialized role, such as endoproteolytic cleavage of VAPB/VPR-1 MSPd in the 20S proteasome as described below.

While the 20S proteasome degrades whole proteins, it can also cleave proteins at precise sites to produce peptides with independent functions. Such examples can be found in the cleaving of translation initiation factors elF3a and eIF4G (61), Y-box RNA/DNA binding protein (62), and Δ40p53 (57). Common across these cleaved peptides is an internal stretch of amino acids known to form a native unfolded or “disordered” structure (61, 62). Both 26S and 20S proteasomes can recognize and internally cleave at this motif (63). Using PONDR Protein Disorder Predictor VLXT (http://www.pondr.com/), we identified a disorder motif downstream the MSPd in VPR-1 (Figure 4). In a separate study, we determined the cleave site of VPR-1 in *C. elegans* to be between Leu156 and Gly157, but no known protease was identified (54). Interestingly, the Leu156, Gly157 cleavage site lies within the strongest and longest predicted disorder motif (Figure 4), suggesting that VAPB/VPR-1 may be processed by the proteasome to release a functional MSPd peptide.

**Figure 4:**
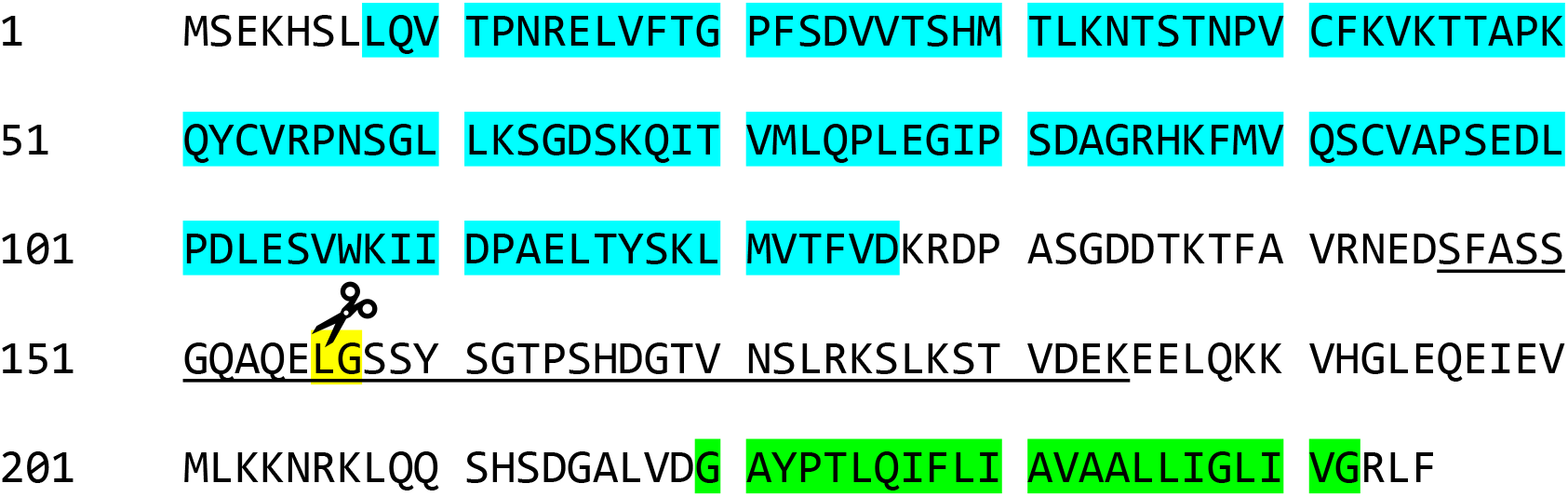
The MSPd cleave site resides in a predicted disordered motif within the VAPB/VPR-1 protein. VPR-1 amino acid sequence was analyzed by PONDR Protein Disorder Predictor VLXT (http://www.pondr.com/) to identify stretches of disordered amino acids. The longest and most likely stretch of disordered amino acids according to the predictor is underlined (aa146-aa184). VPR-1 is cleaved between Leu156 and Gly157 (scissors) to release the MSPd. The MSP domain based on VAPB/VPR-1 MSPd predictions is included the blue highlighted area (11). We predicted the transmembrane domain to be in the highly conserved GXXXG motif (green) (5).

While 20S and 26S proteasomes are usually found unbound in the cytosol, ~40% of 20S proteasomes have been found to be tightly associated with neuronal plasma membranes in mouse (64). These hydrophilic neuronal membrane proteasomes, or NMPs, are exposed to the extracellular space by interacting with GPM6A/B, a multi-pass transmembrane glycoprotein (64). This plasma membrane localization and access to intra- and extra-cellular spaces confers these NMPs the ability to cleave intracellular protein and release the newly generated peptide extracellularly (64). This may also explain how MSPd is cleaved from VAPB/VPR-1 after being trafficked to the plasma membrane by YKT-6 positive vesicles (as discussed above). In fact, neurons are a major cell type that cleaves and secretes MSPd from VAPB/VPR-1 in *Drosophila* and *C. elegans* (12, 13, 15). While this model involving NMPs is appealing, it is unclear at this time whether similar NMPs exist in intestinal or germline cells as they do in neurons, a very specialized cell type. Though our screen is based on the assumption that MSPd is cleaved and secreted via a similar mechanism in all these cell types, each cell lineage may also utilize slightly different mechanisms. Further investigations are needed to identify whether NMPs are found in non-neuronal cells and whether they couple cleavage and release of MSPd from these cells.

### Perspective

This screen aimed to identify players that regulate MSPd cleavage from ER anchored VAPB/VPR-1 and its extracellular secretion. We took an unbiased approach and screened 422 genes in *C. elegans* that were common across the three cell types known to secrete MSPd. One limitation of this screen is that it is based on our understanding of MSPd non-cell-autonomous signaling in *C. elegans*. Mammalian MSPd cleaving and secretion patterns are not known. While various categories of genes came out of this screen, we chose to highlight the v-SNARE (*ykt-6*) and proteasome components (*rpn-1, rpn-3, rpn-10*, and *pbs-2*) due to their previously described roles in the cleavage and secretion of other proteins. As MSPd cleavage and secretion has been implicated in ALS, the results from this screen may have also uncovered new potential players that have not yet been linked to the neurodegenerative disease.

## Acknowledgements

We would like to thank the members of the Miller Lab, Dr. Ekta Tiwary, Dr. Melissa LaBonty, and Dr. Bradley Yoder for their support and valuable discussions regarding this work.

## Funding

This work was funded by the Muscular Dystrophy Association (MDA381893 to M.A.M). Financial training support for H.Z. came from the University of Alabama at Birmingham Translational and Molecular Sciences Pre-doc T32 (GM109780).

## Notes

### Competing Interest Statement

The authors have declared no competing interest.

## References

1. Lev S, Ben Halevy D, Peretti D, Dahan N. The VAP protein family: from cellular functions to motor neuron disease. Trends Cell Biol. 2008;18(6):282–90.

2. Kamemura K, Chihara T. Multiple functions of the ER-resident VAP and its extracellular role in neural development and disease. J Biochem. 2019;165(5):391–400.

3. Weir ML, Klip A, Trimble WS. Identification of a human homologue of the vesicle-associated membrane protein (VAMP)-associated protein of 33 kDa (VAP-33): a broadly expressed protein that binds to VAMP. Biochem J. 1998;333 (Pt 2):247–51.

4. Soussan L, Burakov D, Daniels MP, Toister-Achituv M, Porat A, Yarden Y, et al. ERG30, a VAP-33-related protein, functions in protein transport mediated by COPI vesicles. J Cell Biol. 1999;146(2):301–11.

5. Skehel PA, Fabian-Fine R, Kandel ER. Mouse VAP33 is associated with the endoplasmic reticulum and microtubules. Proc Natl Acad Sci U S A. 2000;97(3):1101–6.

6. Baker AM, Roberts TM, Stewart M. 2.6 A resolution crystal structure of helices of the motile major sperm protein (MSP) of Caenorhabditis elegans. J Mol Biol. 2002;319(2):491–9.

7. Italiano JE, Jr., Roberts TM, Stewart M, Fontana CA. Reconstitution in vitro of the motile apparatus from the amoeboid sperm of Ascaris shows that filament assembly and bundling move membranes. Cell. 1996;84(1):105–14.

8. Wolgemuth CW, Miao L, Vanderlinde O, Roberts T, Oster G. MSP dynamics drives nematode sperm locomotion. Biophys J. 2005;88(4):2462–71.

9. Miller MA, Nguyen VQ, Lee MH, Kosinski M, Schedl T, Caprioli RM, et al. A sperm cytoskeletal protein that signals oocyte meiotic maturation and ovulation. Science. 2001;291(5511):2144–7.

10. Miller MA, Ruest PJ, Kosinski M, Hanks SK, Greenstein D. An Eph receptor sperm-sensing control mechanism for oocyte meiotic maturation in Caenorhabditis elegans. Genes Dev. 2003;17(2):187–200.

11. Han SM, Cottee PA, Miller MA. Sperm and oocyte communication mechanisms controlling C. elegans fertility. Dev Dyn. 2010;239(5):1265–81.

12. Tsuda H, Han SM, Yang Y, Tong C, Lin YQ, Mohan K, et al. The amyotrophic lateral sclerosis 8 protein VAPB is cleaved, secreted, and acts as a ligand for Eph receptors. Cell. 2008;133(6):963–77.

13. Han SM, Tsuda H, Yang Y, Vibbert J, Cottee P, Lee SJ, et al. Secreted VAPB/ALS8 major sperm protein domains modulate mitochondrial localization and morphology via growth cone guidance receptors. Dev Cell. 2012;22(2):348–62.

14. Han SM, El Oussini H, Scekic-Zahirovic J, Vibbert J, Cottee P, Prasain JK, et al. VAPB/ALS8 MSP ligands regulate striated muscle energy metabolism critical for adult survival in caenorhabditis elegans. PLoS Genet. 2013;9(9):e1003738.

15. Cottee PA, Cole T, Schultz J, Hoang HD, Vibbert J, Han SM, et al. The C. elegans VAPB homolog VPR-1 is a permissive signal for gonad development. Development. 2017;144(12):2187–99.

16. Deidda I, Galizzi G, Passantino R, Cascio C, Russo D, Colletti T, et al. Expression of vesicle-associated membrane-protein-associated protein B cleavage products in peripheral blood leukocytes and cerebrospinal fluid of patients with sporadic amyotrophic lateral sclerosis. Eur J Neurol. 2014;21(3):478–85.

17. Nishimura AL, Mitne-Neto M, Silva HC, Richieri-Costa A, Middleton S, Cascio D, et al. A mutation in the vesicle-trafficking protein VAPB causes late-onset spinal muscular atrophy and amyotrophic lateral sclerosis. Am J Hum Genet. 2004;75(5):822–31.

18. Nishimura AL, Mitne-Neto M, Silva HC, Oliveira JR, Vainzof M, Zatz M. A novel locus for late onset amyotrophic lateral sclerosis/motor neurone disease variant at 20q13. J Med Genet. 2004;41(4):315–20.

19. Funke AD, Esser M, Kruttgen A, Weis J, Mitne-Neto M, Lazar M, et al. The p.P56S mutation in the VAPB gene is not due to a single founder: the first European case. Clin Genet. 2010;77(3):302–3.

20. Millecamps S, Salachas F, Cazeneuve C, Gordon P, Bricka B, Camuzat A, et al. SOD1, ANG, VAPB, TARDBP, and FUS mutations in familial amyotrophic lateral sclerosis: genotype-phenotype correlations. J Med Genet. 2010;47(8):554–60.

21. Schultz J, Lee SJ, Cole T, Hoang HD, Vibbert J, Cottee PA, et al. The secreted MSP domain of C. elegans VAPB homolog VPR-1 patterns the adult striated muscle mitochondrial reticulum via SMN-1. Development. 2017;144(12):2175–86.

22. Reinke V, Gil IS, Ward S, Kazmer K. Genome-wide germline-enriched and sex-biased expression profiles in Caenorhabditis elegans. Development. 2004;131(2):311–23.

23. McGhee JD, Sleumer MC, Bilenky M, Wong K, McKay SJ, Goszczynski B, et al. The ELT-2 GATA-factor and the global regulation of transcription in the C. elegans intestine. Dev Biol. 2007;302(2):627–45.

24. Von Stetina SE, Watson JD, Fox RM, Olszewski KL, Spencer WC, Roy PJ, et al. Cell-specific microarray profiling experiments reveal a comprehensive picture of gene expression in the C. elegans nervous system. Genome Biol. 2007;8(7):R135.

25. Brenner S. The genetics of Caenorhabditis elegans. Genetics. 1974;77(1):71–94.

26. Timmons L, Fire A. Specific interference by ingested dsRNA. Nature. 1998;395(6705):854.

27. Kamath RS, Ahringer J. Genome-wide RNAi screening in Caenorhabditis elegans. Methods. 2003;30(4):313–21.

28. Kamath RS, Martinez-Campos M, Zipperlen P, Fraser AG, Ahringer J. Effectiveness of specific RNA-mediated interference through ingested double-stranded RNA in Caenorhabditis elegans. Genome Biol. 2001;2(1):RESEARCH0002.

29. Carim-Todd L, Escarceller M, Estivill X, Sumoy L. Identification and characterization of UBXD1, a novel UBX domain-containing gene on human chromosome 19p13, and its mouse ortholog. Biochim Biophys Acta. 2001;1517(2):298–301.

30. Madsen L, Andersen KM, Prag S, Moos T, Semple CA, Seeger M, et al. Ubxd1 is a novel co-factor of the human p97 ATPase. Int J Biochem Cell Biol. 2008;40(12):2927–42.

31. Ye Y. Diverse functions with a common regulator: ubiquitin takes command of an AAA ATPase. J Struct Biol. 2006;156(1):29–40.

32. Ritz D, Vuk M, Kirchner P, Bug M, Schutz S, Hayer A, et al. Endolysosomal sorting of ubiquitylated caveolin-1 is regulated by VCP and UBXD1 and impaired by VCP disease mutations. Nat Cell Biol. 2011; 13(9):1116–23.

33. Haines DS, Lee JE, Beauparlant SL, Kyle DB, den Besten W, Sweredoski MJ, et al. Protein interaction profiling of the p97 adaptor UBXD1 points to a role for the complex in modulating ERGIC-53 trafficking. Mol Cell Proteomics. 2012;11(6):M111016444.

34. Johnson JO, Mandrioli J, Benatar M, Abramzon Y, Van Deerlin VM, Trojanowski JQ, et al. Exome sequencing reveals VCP mutations as a cause of familial ALS. Neuron. 2010;68(5):857–64.

35. Gkogkas C, Wardrope C, Hannah M, Skehel P. The ALS8-associated mutant VAPB(P56S) is resistant to proteolysis in neurons. J Neurochem. 2011;117(2):286–94.

36. Rossi V, Banfield DK, Vacca M, Dietrich LE, Ungermann C, D’Esposito M, et al. Longins and their longin domains: regulated SNAREs and multifunctional SNARE regulators. Trends Biochem Sci. 2004;29(12):682–8.

37. McNew JA, Sogaard M, Lampen NM, Machida S, Ye RR, Lacomis L, et al. Ykt6p, a prenylated SNARE essential for endoplasmic reticulum-Golgi transport. J Biol Chem. 1997;272(28):17776–83.

38. Zhang T, Hong W. Ykt6 forms a SNARE complex with syntaxin 5, GS28, and Bet1 and participates in a late stage in endoplasmic reticulum-Golgi transport. J Biol Chem. 2001;276(29):27480–7.

39. Fukasawa M, Varlamov O, Eng WS, Sollner TH, Rothman JE. Localization and activity of the SNARE Ykt6 determined by its regulatory domain and palmitoylation. Proc Natl Acad Sci U S A. 2004;101(14):4815–20.

40. Dietrich LE, Gurezka R, Veit M, Ungermann C. The SNARE Ykt6 mediates protein palmitoylation during an early stage of homotypic vacuole fusion. EMBO J. 2004;23(1):45–53.

41. Hasegawa H, Zinsser S, Rhee Y, Vik-Mo EO, Davanger S, Hay JC. Mammalian ykt6 is a neuronal SNARE targeted to a specialized compartment by its profilin-like amino terminal domain. Mol Biol Cell. 2003;14(2):698–720.

42. Meiringer CT, Auffarth K, Hou H, Ungermann C. Depalmitoylation of Ykt6 prevents its entry into the multivesicular body pathway. Traffic. 2008;9(9):1510–21.

43. Kriegenburg F, Bas L, Gao J, Ungermann C, Kraft C. The multi-functional SNARE protein Ykt6 in autophagosomal fusion processes. Cell Cycle. 2019;18(6-7):639–51.

44. Tochio H, Tsui MM, Banfield DK, Zhang M. An autoinhibitory mechanism for nonsyntaxin SNARE proteins revealed by the structure of Ykt6p. Science. 2001;293(5530):698–702.

45. Pylypenko O, Schonichen A, Ludwig D, Ungermann C, Goody RS, Rak A, et al. Farnesylation of the SNARE protein Ykt6 increases its stability and helical folding. J Mol Biol. 2008;377(5):1334–45.

46. Ungermann C, von Mollard GF, Jensen ON, Margolis N, Stevens TH, Wickner W. Three v-SNAREs and two t-SNAREs, present in a pentameric cis-SNARE complex on isolated vacuoles, are essential for homotypic fusion. J Cell Biol. 1999;145(7):1435–42.

47. Kweon Y, Rothe A, Conibear E, Stevens TH. Ykt6p is a multifunctional yeast R-SNARE that is required for multiple membrane transport pathways to the vacuole. Mol Biol Cell. 2003;14(5):1868–81.

48. Tai G, Lu L, Wang TL, Tang BL, Goud B, Johannes L, et al. Participation of the syntaxin 5/Ykt6/GS28/GS15 SNARE complex in transport from the early/recycling endosome to the trans-Golgi network. Mol Biol Cell. 2004;15(9):4011–22.

49. Nair U, Jotwani A, Geng J, Gammoh N, Richerson D, Yen WL, et al. SNARE proteins are required for macroautophagy. Cell. 2011;146(2):290–302.

50. Gordon DE, Chia J, Jayawardena K, Antrobus R, Bard F, Peden AA. VAMP3/Syb and YKT6 are required for the fusion of constitutive secretory carriers with the plasma membrane. PLoS Genet. 2017;13(4):e1006698.

51. Gao J, Reggiori F, Ungermann C. A novel in vitro assay reveals SNARE topology and the role of Ykt6 in autophagosome fusion with vacuoles. J Cell Biol. 2018;217(10):3670–82.

52. Takats S, Glatz G, Szenci G, Boda A, Horvath GV, Hegedus K, et al. Non-canonical role of the SNARE protein Ykt6 in autophagosome-lysosome fusion. PLoS Genet. 2018;14(4):e1007359.

53. Gross JC, Chaudhary V, Bartscherer K, Boutros M. Active Wnt proteins are secreted on exosomes. Nat Cell Biol. 2012;14(10):1036–45.

54. Zein-Sabatto H, Cole T, Hoang HD, Tiwary E, Chang C, Miller MA. The type II integral ER membrane protein VAP-B homolog in C. elegans is cleaved to release the N-terminal MSP domain to signal non-cell-autonomously. Dev Biol. 2020;470:10–20.

55. Morozov AV, Karpov VL. Biological consequences of structural and functional proteasome diversity. Heliyon. 2018;4(10):e00894.

56. Papaevgeniou N, Chondrogianni N. The ubiquitin proteasome system in Caenorhabditis elegans and its regulation. Redox Biol. 2014;2:333–47.

57. Olshina MA, Ben-Nissan G, Sharon M. Functional regulation of proteins by 20S proteasome proteolytic processing. Cell Cycle. 2018;17(4):393–4.

58. Ehlinger A, Walters KJ. Structural insights into proteasome activation by the 19S regulatory particle. Biochemistry. 2013;52(21):3618–28.

59. Young P, Deveraux Q, Beal RE, Pickart CM, Rechsteiner M. Characterization of two polyubiquitin binding sites in the 26 S protease subunit 5a. J Biol Chem. 1998;273(10):5461–7.

60. Shimada M, Kanematsu K, Tanaka K, Yokosawa H, Kawahara H. Proteasomal ubiquitin receptor RPN-10 controls sex determination in Caenorhabditis elegans. Mol Biol Cell. 2006;17(12):5356–71.

61. Baugh JM, Pilipenko EV. 20S proteasome differentially alters translation of different mRNAs via the cleavage of eIF4F and eIF3. Mol Cell. 2004;16(4):575–86.

62. Sorokin AV, Selyutina AA, Skabkin MA, Guryanov SG, Nazimov IV, Richard C, et al. Proteasome-mediated cleavage of the Y-box-binding protein 1 is linked to DNA-damage stress response. EMBO J. 2005;24(20):3602–12.

63. Liu CW, Corboy MJ, DeMartino GN, Thomas PJ. Endoproteolytic activity of the proteasome. Science. 2003;299(5605):408–11.

64. Ramachandran KV, Margolis SS. A mammalian nervous-system-specific plasma membrane proteasome complex that modulates neuronal function. Nat Struct Mol Biol. 2017;24(4):419–30.

